# Translational plasticity facilitates the accumulation of nonsense genetic variants in the human population

**DOI:** 10.1101/038687

**Authors:** Sujatha Jagannathan, Robert K. Bradley

## Abstract

Genetic variants that disrupt protein-coding DNA are ubiquitous in the human population, with ~100 such loss-of-function variants per individual. While most loss-of-function variants are rare, a subset have risen to high frequency and occur in a homozygous state in healthy individuals. It is unknown why these common variants are well-tolerated, even though some affect essential genes implicated in Mendelian disease. Here, we combine genomic, proteomic, and biochemical data to demonstrate that many common nonsense variants do not ablate protein production from their host genes. We provide computational and experimental evidence for diverse mechanisms of gene rescue, including alternative splicing, stop codon readthrough, alternative translation initiation, and C-terminal truncation. Our results suggest a molecular explanation for the mild fitness costs of many common nonsense variants, and indicate that translational plasticity plays a prominent role in shaping human genetic diversity.

## INTRODUCTION

The discovery of pervasive genetic variation that is predicted to disrupt protein-coding DNA sequences is a surprising finding of recent human genetics studies. This loss-of-function genetic variation can arise from many sources, including single nucleotide variants (SNVs) that introduce stop codons, insertions or deletions that disrupt the reading frame, disruption of splice sites, or large structural variation. While loss-of-function variants are generally found at low allelic frequencies relative to synonymous variants (**Figure 1A**), many have risen to sufficiently high frequency to be present in a homozygous state. A careful study of these variants estimated that typical individuals carry ~100 loss-of-function variants, of which ~20 are present in a homozygous state (MacArthur et al. 2012).

**Figure 1.**
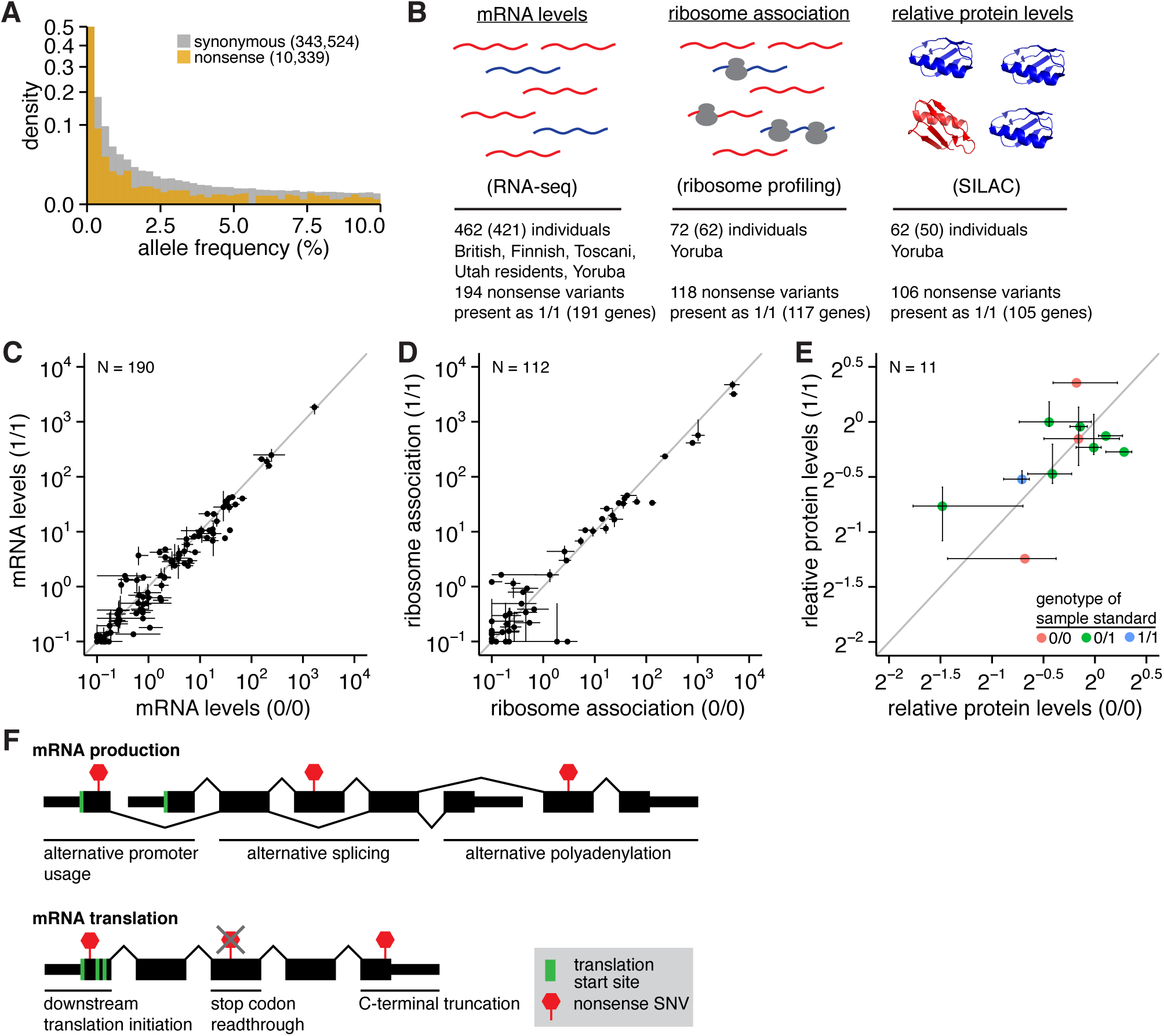
Putatively disabled genes exhibit normal levels of the encoded proteins. (**A**) Histogram of alternate allele frequencies for synonymous and nonsense variants identified by the 1000 Genomes Project (1000 Genomes Project Consortium 2012). (**B**) Genome-wide measurements of mRNA abundance (Lappalainen et al. 2013), mRNA:ribosome association (Battle et al. 2015), and protein abundance (Battle et al. 2015) in lymphoblastoid cell lines (LCLs) that we analyzed here. This data was available for 462, 72, and 62 individuals, respectively; however, we restricted our analyses to the 421, 62, and 50 individuals that were genotyped by sequencing. The represented populations are British in England and Scotland, Finnish in Finland, Toscani in Italy, Utah residents with Northern and Western European ancestry, and Yoruba in Ibadan, Nigeria. SILAC, stable isotope labeling by amino acids in cell culture (Ong et al. 2002). (**C**) Median RNA levels of CDSs containing nonsense variants, averaged across samples that are homozygous for the reference (0/0) or alternate (1/1) alleles. Error bars, first and fourth quartiles of expression across all 0/0 or 1/1 samples. Units, fragments per kilobase per million mapped reads (FPKM) of the CDS containing each variant. N, number of analyzed variants. Analysis based on the 194 nonsense variants stated in (B), but restricted to variants within Ensembl coding genes that were present as both 0/0 and 1/1 in samples with RNA-seq data. (**D**) As (C), but illustrates mRNA:ribosome association as measured by ribosome profiling. Units, fragments per kilobase per million mapped reads (FPKM) of the CDS containing each variant. Analysis based on the 118 nonsense variants stated in (B), but restricted to variants within Ensembl coding genes that were present as both 0/0 and 1/1 in samples with ribosome profiling data. (**E**) As (C), but illustrates relative levels of proteins encoded by CDSs containing nonsense variants as measured by SILAC mass spectrometry. Units, protein levels relative to the sample standard (e.g., 2^0^ indicates no change in protein levels relative to the standard), whose genotype for each variant is indicated by the point color. Protein levels were taken directly from Battle et al, who estimated protein levels as the median sample:standard ratio across all peptides arising from a single parent gene in each sample (Battle et al. 2015). Analysis based on the 106 nonsense variants stated in (B), but restricted to variants within Ensembl coding genes that were present as both 0/0 and 1/1 in samples with SILAC data. Fewer variants can be analyzed here than in (C-D) due to the low coverage of mass spectrometry data relative to RNA-seq or ribosome profiling. (**F**) Possible mechanisms to enable protein production from genes containing nonsense variants. Nonsense variants may be isoform-specific, result in N- or C-terminal truncation of the encoded protein, or be subject to readthrough during translation.

While deleterious genetic variants can rise to high frequencies in specific populations due to demographic effects such as population bottlenecks (Lim et al. 2014), the high allele frequencies of some loss-of-function variants within diverse and large human populations suggests that these variants are subject to relatively weak purifying selection. In some cases, gene inactivation may confer a fitness benefit for individuals or the species, as has been posited for *CASP12* (Xue et al. 2006), *ACTN3* (MacArthur et al. 2007), and *ERAP2* (Andrés et al. 2010), and positive or balancing selection may drive the inactive allele to high frequency. However, the rarity of most loss-of-function variants relative to synonymous variants suggests that beneficial loss-of-function variants are exceptions rather than the norm.

Consistent with the apparently mild purifying selection acting on common loss-of-function variants, a recent study of five European populations reported that no homozygous loss-of-function variants within these populations were associated with detectable phenotypic deviance (Kaiser et al. 2015). This report is consistent with expectations based on a gene disruption screen in zebrafish, which found that only 6% of randomly induced nonsense or splice site mutations yielded an embryonic phenotype (Kettleborough et al. 2013). Knockout studies in mouse models yielded much higher estimates of mouse embryonic lethality (Ayadi et al. 2012; de Angelis et al. 2015). However, it is difficult to directly compare these studies, particularly because gene inactivation in the murine models was frequently achieved by unambiguously disrupting the coding DNA via insertion of a polyadenylation site within the gene body (versus the more modest genetic changes induced by *N*-ethyl-*N*-nitrosourea (ENU) mutagenesis in the zebrafish models).

The apparently mild fitness consequences of common loss-of-function variants can be explained in two different ways, depending upon whether these variants affect non-essential or essential genes. First, genetic redundancy may render particular genes dispensable, such that they can be inactivated without significant fitness costs. Loss-of-function variants are enriched in genes with more paralogs than other genes, most notably the olfactory receptors, as well as genes that are relatively poorly conserved (Gilad et al. 2003; MacArthur et al. 2012). Second, loss-of-function variants may disrupt coding DNA yet not completely disable their (essential) host genes’ functions. Consistent with this hypothesis, previous studies noted that many loss-of-function variants do not affect all isoforms of their parent genes, and furthermore that loss-of-function variants are enriched near the 3’ ends of coding sequences (MacArthur et al. 2012; Liu and Lin 2015; Kaiser et al. 2015). These observations suggest that some loss-of-function variants may not completely destroy the coding potential of their host genes, potentially explaining why those variants are well-tolerated. However, this hypothesis—that many loss-of-function variants do not ablate their host genes’ functions—has not been experimentally tested.

Here, we systematically determined the consequences of common nonsense variants for mRNA translation and protein production using a combination of genome-wide data and directed biochemical experiments.

## RESULTS

### Many common nonsense variants do not disrupt protein production

We set out to test the hypothesis that many common loss-of-function variants do not ablate protein production from their host genes, thereby explaining the apparently mild phenotypic consequences of these variants even when found in a homozygous state. To test this hypothesis, we took advantage of several recently published datasets representing successive stages of gene expression. These datasets consist of genome-wide measurements of mRNA levels by RNA-seq (Lappalainen et al. 2013), mRNA:ribosome association by ribosome profiling (Battle et al. 2015), and protein abundance by quantitative mass spectrometry (Battle et al. 2015) in lymphoblastoid cell lines (LCLs) that were established by the HapMap and 1000 Genomes Projects (International HapMap 3 Consortium 2010; 1000 Genomes Project Consortium 2012). For each dataset, we used published sample genotypes to identify single nucleotide variants (SNVs) that induced synonymous or nonsense codon changes with respect to the reference genome in one or more RefSeq coding transcripts.

We specifically focused our analysis on common nonsense variants that were present in a homozygous state and thereby predicted to induce complete gene knockouts. We focused on nonsense variants rather than the broader spectrum of loss-of-function variation because nonsense variants can be identified in a straightforward manner from both DNA and RNA sequencing, their impact upon the mRNA is unambiguous, they are relevant to many Mendelian diseases, and their allelic expression can be reliably quantified from RNA-seq data. Of all nonsense variants that we identified, a total of 194 (RNA-seq), 118 (ribosome profiling), and 106 (mass spectrometry) nonsense variants were present in a homozygous state in the genome of at least one assayed sample in each respective dataset (**Figure 1B**).

By studying individuals with either two (genotype 0/0; 0 = reference allele) or zero (genotype 1/1; 1 = alternate allele) intact gene copies, we were able to unambiguously determine how each nonsense variant affected protein production. Restricting to those variants, we found that levels of the parent mRNAs containing each nonsense variant were similar in 0/0 and 1/1 individuals (Figure 1C). The similarity of mRNA levels in 0/0 and 1/1 individuals suggested that many of the mRNAs containing the nonsense variants studied here escaped from nonsense-mediated decay (NMD). To determine the effect of NMD on these variants, we quantified allele-specific expression (ASE) of the alternate alleles illustrated in **Figure 1C** in heterozygous (genotype 0/1) individuals. We restricted our analysis to nonsense variants covered by at least 20 RNA-seq reads in at least one heterozygous sample (so that we could accurately measure ASE) and computed the median ASE across all such heterozygous samples for each variant. The 36 nonsense variants meeting this coverage requirement exhibited a median ASE of 45%, only modestly lower than the 50% expected if both alleles were equally transcribed and not affected by NMD. Seven of the 36 variants exhibited a median ASE < 10%, consistent with relatively efficient degradation by NMD (**Figure S1A**). To test whether the coverage threshold of 20 reads biased our conclusions, we repeated the analysis using thresholds of 5, 10, 20, and 50 reads. Those thresholds yielded median ASE estimates of 40%, 41%, 45%, and 45% for nonsense variants (N = 59, 52, 36, and 25 variants analyzed), indicating that our results are robust with respect to the specific read threshold used.

We next tested whether nonsense variants that efficiently triggered NMD were predicted to do so. Stop codons that lie ≥50 nt upstream of exon-exon junctions are canonically predicted to trigger NMD (Popp and Maquat 2013; Lykke-Andersen and Jensen 2015). 86% of the seven variants with ASE < 10% were predicted NMD substrates by the “50 nt” rule, while 62% of the 29 variants with ASE ≥10% were predicted NMD substrates. We conclude that the majority of mRNAs containing the nonsense variants studied here escape NMD, although a subset are efficiently degraded, consistent with previous reports (MacArthur et al. 2012; Kukurba et al. 2014; Rivas et al. 2015).

The observed similarity in levels of 0/0 and 1/1 mRNAs extended to mRNA translation as well. Association between the parent mRNAs and ribosomes, as well as levels of the encoded proteins, was similar between 0/0 and 1/1 individuals (**Figure 1D-E**). (The much lower coverage of both ribosome profiling and mass spectrometry data relative to RNA-seq data rendered the analysis of allele-specific expression in 0/1 individuals infeasible for these assays of mRNA translation.) We observed no cases where homozygous nonsense variants completely abolished mRNA expression, mRNA:ribosome association, or protein expression, with the caveat that these measurements were necessarily restricted to variants within genes that were detectably expressed in LCLs by RNA-seq, ribosome profiling, or mass spectrometry.

Genotyping errors could potentially explain why we observed normal protein levels from genes containing homozygous nonsense variants. We therefore used RNA-seq to identify and remove incorrectly called genotypes. For each nonsense variant, we restricted our analyses to samples for which the variant was covered by at least ten reads, and removed samples that were annotated as 0/0 or 1/1 but exhibited expression of the alternate or reference alleles, respectively. While we did identify and remove a small number of such variant-sample pairs whose genotypes were inconsistent with the observed allelic expression, removing those inconsistent data points did not affect our conclusion that neither mRNA expression, mRNA:ribosome association, nor protein expression was abolished by the analyzed homozygous nonsense variants (**Figure S1B-D**). Therefore, genotyping errors do not explain why these nonsense variants do not abolish protein production.

### Permissive RNA processing may enable protein production from genes containing nonsense variants

The extensive regulation that occurs between the initiation of gene transcription and the completion of mRNA translation offers ample opportunities for protein production from genes containing nonsense variants (although gene function may or may not be preserved). Protein can be produced from genes containing nonsense variants by one of two means (**Figure 1F**). First, a nonsense variant may be isoform-specific, such that at least one coding isoform of the parent gene does not contain the variant. In this case, the nonsense variant may prevent translation of one or more, but not all, isoforms. Second, the stop codon introduced by a nonsense variant may not be sufficient to abolish productive translation. This can occur if the ribosome can initiate translation downstream of the stop codon, if the stop codon is subject to readthrough, or if the nonsense variant is near the normal stop codon.

Sequence analyses reported in previous studies suggest that both alternative splicing and protein truncation may contribute to the robust protein production from genes containing nonsense variants that we observed (**Figure 1C-E**). Approximately one-third of previously studied nonsense variants are isoform-specific (MacArthur et al. 2012; Kaiser et al. 2015); minor coding isoforms are enriched for nonsense variants relative to major isoforms (Liu and Lin 2015); nonsense variants preferentially occur at the 3’ ends of their host CDSs (MacArthur et al. 2012; Sulem et al. 2015). We therefore set out to systematically identify molecular mechanisms that enabled protein production from genes containing nonsense variants. Our general strategy was to combine RNA-seq, ribosome profiling, and SILAC data to identify the mechanisms that enabled protein production from genes containing the common nonsense variants studied in **Figure 1**, and then computationally predict the likely relevance of each mechanism to lower-frequency nonsense variants using sequence analysis of the ~10,000 nonsense variants identified by the 1000 Genomes Project.

### Nonsense variants are frequently isoform-specific and removed by alternative splicing

We first tested whether nonsense variants are more frequently isoform-specific than their synonymous counterparts. For each nonsense variant within a multi-exon gene, we determined whether the gene contained at least one RefSeq coding transcript for which the alternate allele did not introduce an in-frame stop codon, or if the nonsense variant was contained within alternatively spliced sequence of the mRNA. We similarly measured isoform specificity for synonymous variants, restricted to the subset of synonymous variants lying within genes that contain nonsense variants in order to control for the fact that nonsense variants preferentially occur in specific gene classes (MacArthur et al. 2012; Sulem et al. 2015). We eliminated variants lying within genes encoding olfactory receptors from this and subsequent analyses, as this large gene family has been subject to frequent and likely largely neutral gene inactivation during human evolution (Gilad et al. 2003).

Approximately 28% of synonymous variants in multi-exon genes are isoform-specific. In order to test for a relationship between isoform specificity and the prevalence of variants, we classified each variant as rare, low-frequency, or common, corresponding to allele frequencies of ≤ 0.5%, 0.5-5%, and ≥5% in the 1000 Genomes data. We observed no relationship between isoform specificity of synonymous variants and their frequency in the human population, consistent with our expectation that most synonymous codon substitutions are functionally neutral. In contrast, approximately 31% of rare nonsense variants are isoform-specific, rising to 39% and 43% of low-frequency and common nonsense variants, suggesting that isoform-specific nonsense variants are subject to relaxed purifying selection (**Figure 2A**). When we considered all synonymous variants rather than restricting to those within genes containing nonsense variants, we observed slightly greater disparities between the isoform specificity of synonymous versus nonsense variants (**Figure S2A**). Our estimates of isoform specificity for nonsense variants are consistent with a previous estimate of 32.3% based on a smaller cohort of 185 individuals (MacArthur et al. 2012).

**Figure 2.**
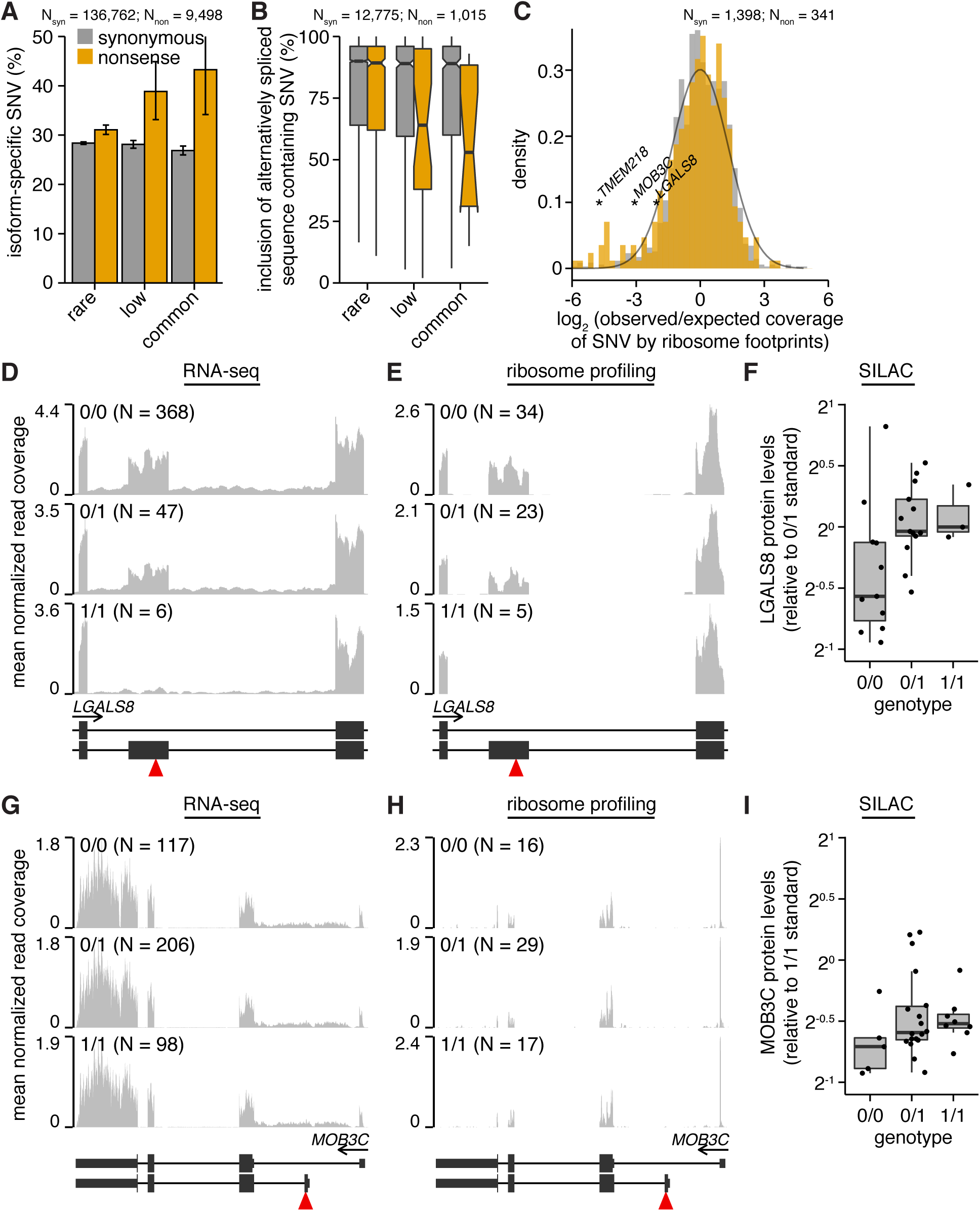
Alternative splicing, promoter usage, and polyadenylation remove nonsense genetic variants from mature mRNA. (**A**) Percentages of variants that are isoform-specific. Isoform-specific variants are defined as those that do not induce the indicated coding change within at least one RefSeq CDS of the parent gene or that lie within known alternatively spliced mRNA sequence. Plot restricted to variants that lie within genes containing nonsense variants. N, number of analyzed variants. Error bars, 95% confidence intervals as estimated by the binomial proportion test. (**B**) Median inclusion of variants lying within known alternatively spliced mRNA sequence across 16 human tissues. An inclusion level of 75% indicates that 75% of mRNAs transcribed from the parent gene contain the variant, while 25% do not. Inclusion was computed using the Body Map 2.0 data. Notches, approximate 95% confidence interval for the median. Plot restricted to variants that lie within genes containing nonsense variants. (**C**) Histogram of observed versus expected coverage of variants by ribosome footprints. Observed coverage was computed as the number of footprints overlapping each variant; expected coverage was computed as the total number of footprints overlapping each variant’s host CDS, normalized such that the median of the ratio observed:expected over all variants was equal to 1. All footprint coverage calculations were restricted to 0/0 samples to avoid potentially confounding effects of nonsense variants, and the plotted values indicate medians over those samples. Black line, best-fit normal distribution estimated from synonymous variants. Plot restricted to variants that lie within genes containing nonsense variants and that are present in samples for which ribosome profiling data was available. (**D**) RNA-seq and (**E**) ribosome profiling read coverage of a nonsense variant lying within a cassette exon of *LGALS8*, stratified by sample genotype. Coverage was normalized per sample to control for sequencing depth and then averaged over all samples with the indicated genotypes. Units, reads per million. Red triangle, location of nonsense variant. (**F**) LGALS8 protein levels relative to a sample standard with genotype 0/1. (**G**) RNA-seq and (**H**) ribosome profiling read coverage of a nonsense variant lying within an alternate 5’ exon of *MOB3C*. (**I**) MOB3C protein levels relative to a sample standard with genotype 0/1.

The functional consequences of an isoform-specific variant depend upon the frequency with which those isoforms are produced as mature mRNAs. To test whether nonsense variants preferentially fall within isoforms that are used infrequently, we quantified the inclusion of alternatively spliced sequence containing synonymous or nonsense variants across sixteen human tissues. Isoform-specific synonymous variants exhibited a median inclusion of ~90% irrespective of allele frequency, while low-frequency and common isoform-specific nonsense variants exhibited median inclusions of 64% and 53% (Figure 2B, S2B). We conclude that even among the isoform-specific nonsense variants, those variants that are most frequently spliced out of mature mRNA are subject to further relaxed selection.

We next measured the empirical isoform specificity of the specific set of homozygous nonsense variants that we analyzed in LCLs (**Figure 1C-E**). For each SNV within an expressed gene, we quantified the expected versus observed number of ribosome footprints overlapping the SNV as a simple empirical measure of isoform specificity (e.g., an isoform-specific SNV is expected to exhibit a ratio observed:expected less than 1). We estimated the expected footprint coverage from the total number of footprints aligned to the parent transcript of each SNV and averaged this footprint coverage over all samples with a 0/0 genotype for that SNV. (We restricted the coverage analysis to 0/0 samples in order to measure position-specific mRNA:ribosome association in the absence of premature stop codons.) The resulting distributions were similar for most synonymous and nonsense variants, consistent with our genome-wide prediction that the majority of variants are not isoform-specific. Nonetheless, a subset of nonsense variants were strongly depleted for ribosome footprint coverage beyond the background distribution estimated from synonymous variants, indicating that they are frequently excluded from mature mRNAs engaged with ribosomes in LCLs (**Figure 2C**). Low-frequency and common nonsense variants exhibited greater depletion for footprint coverage relative to rare nonsense variants, again consistent with relaxed purifying selection (**Figure 2C**).

Each of the mechanisms proposed in **Figure 1F** contributed to the isoform specificity of nonsense variants. Specific examples include nonsense variants within a frame-preserving cassette exon of *LGALS8*, an alternate promoter of *MOB3C*, and alternatively spliced sequence of *TMEM218* that is either coding or non-coding, depending upon which start codon is used (**Figure 2D-I**, **S2D-G**). SILAC data was available for LGALS8 and MOB3C, allowing us to confirm that protein levels were sustained in LCLs with homozygous copies of the corresponding nonsense variants. In both cases, protein levels were higher for 0/1 and 1/1 samples relative to 0/0 samples, perhaps due to compensatory translational up-regulation in individuals carrying the alternate alleles or differential stability of the relevant protein isoforms (**Figure 2F, I**).

As alternative splicing is frequently tissue-specific (Wang et al. 2008), the degree to which isoform specificity mitigates the consequences of a nonsense variant may differ substantially between different cell types. For example, the *LGALS8* cassette exon is highly tissue-specific, with inclusion ranging from 5% in colon to 87% in testes of individuals with genotype 0/0 (inclusion is ~55% in LCLs with genotype 0/0; **Figure S2E**). Therefore, both the levels of LGALS8 protein in 1/1 individuals and the consequences of specifically ablating the inclusion isoform may differ between cell types.

### Nonsense variants may be subject to stop codon readthrough

Genes containing nonsense variants within constitutively included sequence cannot produce mRNAs lacking the variants, but could potentially produce protein nonetheless through permissive mRNA translation (**Figure 1F**). We therefore searched for read coverage patterns indicative of potential stop codon readthrough, wherein the translating ribosome does not efficiently terminate at the stop codon, but rather decodes the stop codon as a sense codon and continues elongating. A nonsense variant that is subject to readthrough might suppress levels of the encoded protein due to imperfectly efficient readthrough, but would induce only a single amino acid change at the nonsense variant.

To identify potential examples of readthrough, we compared ribosome footprint coverage of genes containing nonsense variants in samples with genotypes 0/0 or 1/1 for the variants. Of the 118 nonsense variants present as 1/1 in at least one sample within the Yoruba cohort (**Figure 1B**), 15 variants had both 0/0 and 1/1 samples available and lay within genes with sufficient footprint coverage in LCLs to identify potential readthrough (median coverage of relevant transcripts of ~1 footprint per million mapped reads). Of those 15, two nonsense variants within *PVRIG* and *SLFN13* exhibited similar patterns of mRNA:ribosome association in 0/0, 0/1, and 1/1 samples, including in the immediately vicinity of the premature termination codon (**Figure 3**, **S3**). This pattern of ribosome footprints is consistent with stop codon readthrough, in the sense that there is similar quantitative read coverage before and after the nonsense variant. However, it is important to note that this pattern of read coverage could potentially arise from ribosome scanning and translation initiation downstream of the nonsense variant as well.

**Figure 3.**
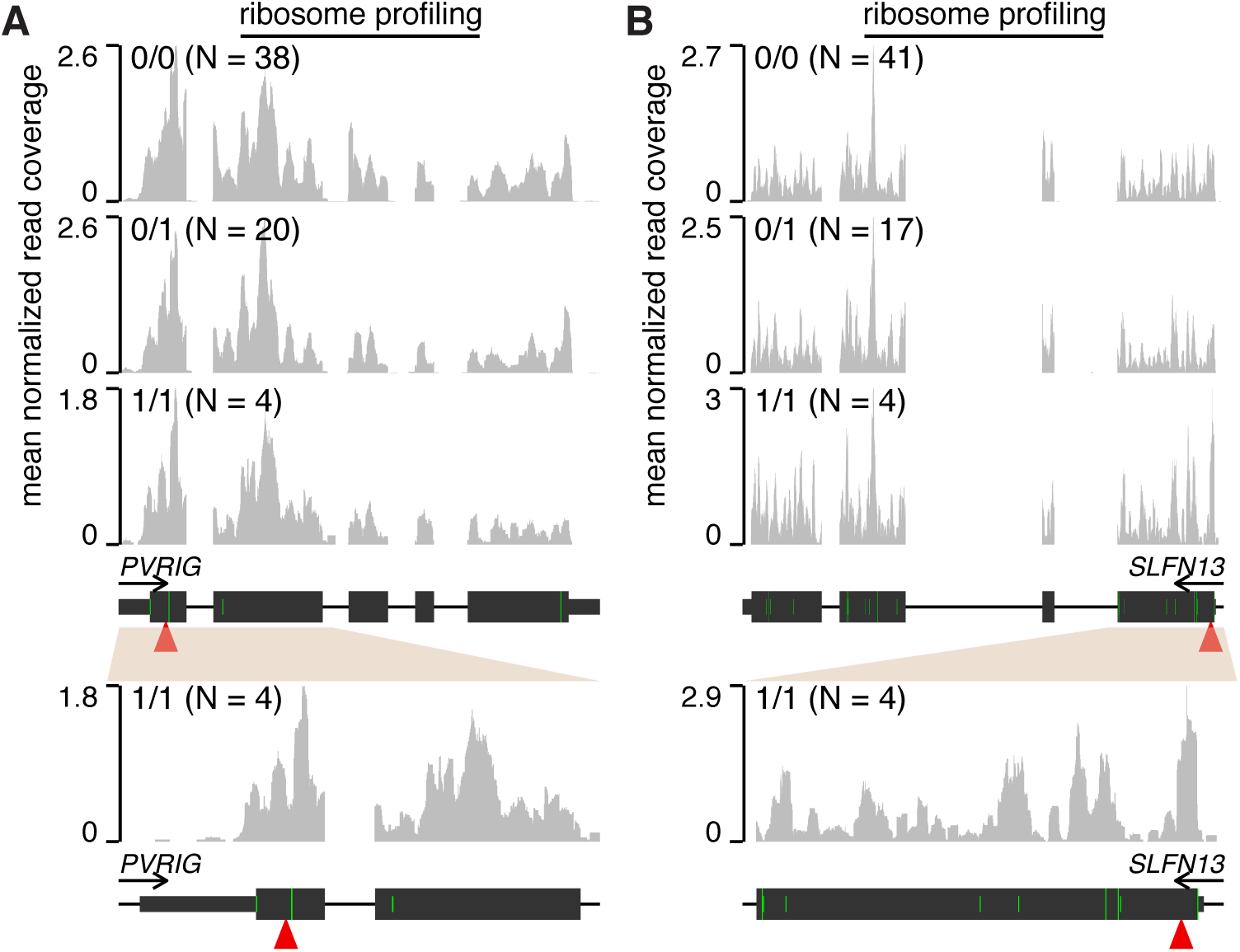
Stop codon readthrough likely enables translation of nonsense variant-containing mRNAs. (**A**) Ribosome profiling read coverage of nonsense variants lying within *PVRIG* and (**B**) *SLFN13*. Units and data normalization are as in Figure 2. Bottom plots are zoomed-in versions of the top plots. Green bars, ATG start codons. Full/half-height green bars indicate ATG codons that are/are not within a Kozak consensus context, defined as RnnATGG (R = A/G).

We tested whether our analysis of *PVRIG* and *SLFN13* was confounded by mis-mapping of ribosome footprints from homologous transcribed genomic loci. The first two coding exons of *PVRIG*, which contain the nonsense variant, have 96.8% homology to the transcribed pseudogene *PVRIG2P*. However, as the density of mismatches between *PVRIG* and *PVRIG2P* is high enough to uniquely map ribosome profiling reads and *PVRIG2P* exhibits no ribosome footprint coverage, we conclude that the ribosome association illustrated in **Figure 3A** likely arises from the PVRIG mRNA and not a homologous locus. Similarly, while the first coding exon of *SLFN13*, which contains the nonsense variant, has 82.9% homology to the first coding exon of the expressed gene *SLFN11*, the density of mismatches is sufficient to uniquely map ribosome footprints. Finally, both RNA-seq and ribosome profiling reads mapping to the *PVRIG* and *SLFN13* nonsense variants supported 100% expression of the alternate allele in 1/1 samples, indicating that genotyping errors are unlikely.

We conclude that stop codon readthrough may enable translation of some mRNAs containing nonsense variants. However, we were unable to estimate the fraction of nonsense variants that might be so rescued given the infeasibility of predicting readthrough from mRNA sequence alone.

### Nonsense variants frequently induce N- or C-terminal protein truncation

In addition to stop codon readthrough, stable expression of a truncated protein might enable protein production from a nonsense variant-containing mRNA. Protein truncation can occur at the N or C terminus through different mechanisms. An N-terminally truncated protein results from translation initiation at a start site downstream of the nonsense variant. In contrast, C-terminal truncation results if the nonsense variant is sufficiently close to the normal stop codon and the host transcript is not efficiently degraded via NMD.

We took a sequence-based approach to identify nonsense variants that might induce N- or C-terminal truncation. Simply plotting the relative positions of nonsense variants within their host CDSs revealed that these variants are enriched at the 3’ ends of their host CDSs and otherwise uniformly distributed, while synonymous variants exhibited no peak at the 3’ end (**Figure 4A**, **S4A**). This pattern is consistent with previous studies of different cohorts (MacArthur et al. 2012; Sulem et al. 2015) and suggests that C-terminal truncation occurs frequently. We did not observe any enrichment for nonsense variants at the 5’ ends of their host CDSs, suggesting that start codon mis-annotation—which can cause a variant within a 5’ untranslated region to be incorrectly labeled as a nonsense variant (MacArthur and Tyler-Smith 2010; Sulem et al. 2015)—is not a frequent confounding factor in our analyses.

**Figure 4.**
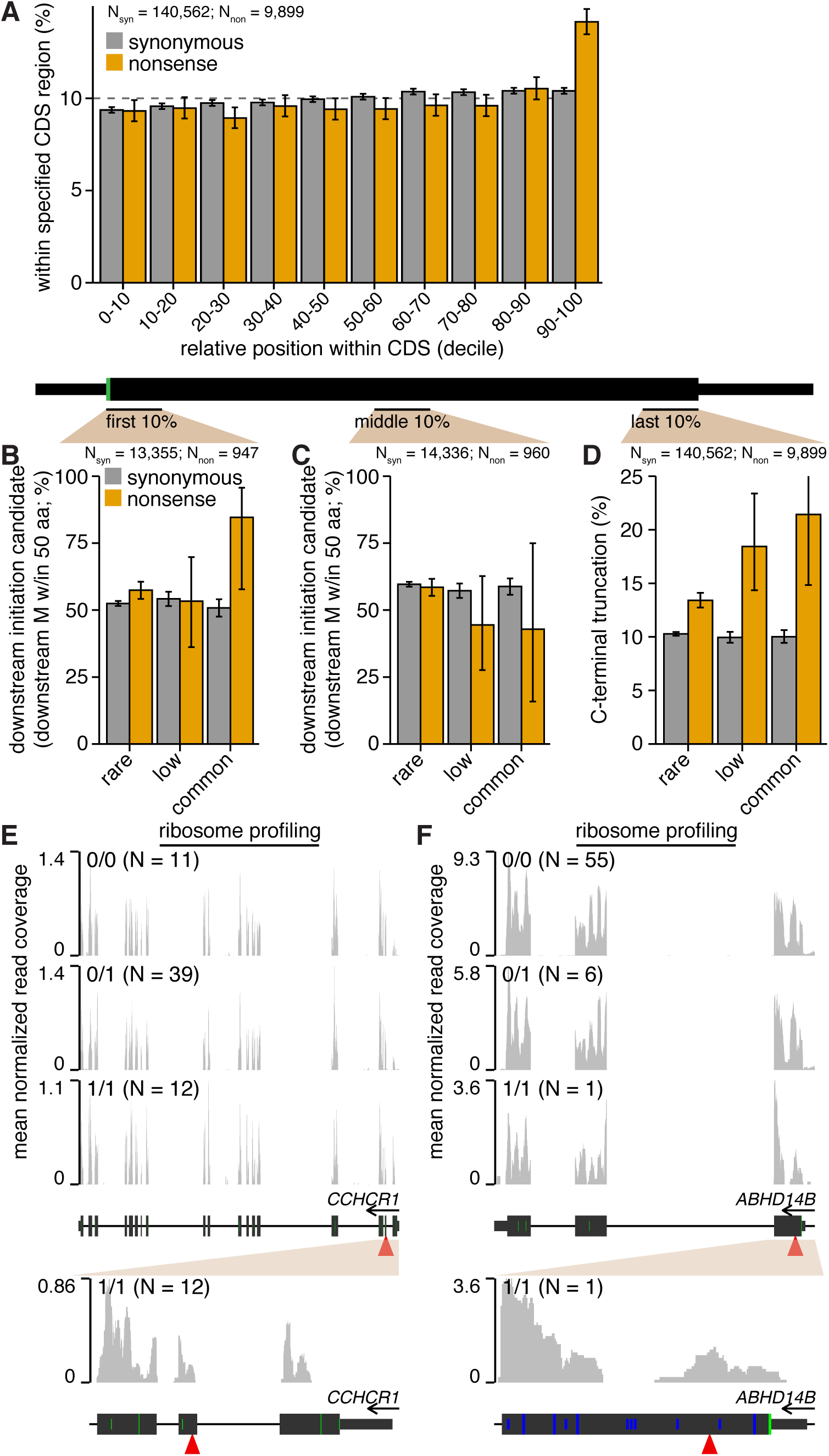
N- and C-terminal protein truncation enable translation of nonsense variant-containing mRNAs. (**A**) Positional distribution of variants within their host transcripts. Plot indicates the percentages of variants lying within the indicated deciles of their host CDSs’ lengths. Error bars, 95% confidence intervals as estimated by the binomial proportion test. (**B**) Percentages of variants lying within the first 10% or (**C**) middle 10% of their host CDS’s length that have a downstream methionine (M) within 50 amino acids. Plot restricted to variants that lie within genes containing nonsense variants. Error bars, 95% confidence intervals as estimated by the binomial proportion test. (**D**) Percentages of variants lying within the last 10% of their host CDS’s length. Plot restricted to variants that lie within genes containing nonsense variants. (**E**) Ribosome profiling read coverage of nonsense variants lying within *CCHCR1* and (**F**) *ABHD14B*. Units, data normalization, and notation are as in Figure 3. Blue bars, CTG or GTG non-canonical start codons (no downstream ATG codons are present within the plot region). Full/half-height blue bars indicate CTG or GTG codons that are/are not within a Kozak consensus context, defined as RnnSTGG (R = A/G, S = C/G).

To test whether N-terminal truncation via translation initiation at downstream start sites might also occur, we restricted to SNVs lying within the first 10% of their host CDSs and tested whether those variants were followed by an in-frame methionine within 50 amino acids downstream of the SNV. Approximately 52% of such synonymous variants had a downstream methionine within this distance. In contrast, both rare and common nonsense variants exhibited statistically significant enrichment for downstream methionines, with 57% and 85% of such variants categorized as potential candidates for downstream initiation (**Figure 4B**). We next performed two controls to confirm that methionines were specifically enriched downstream of nonsense variants that were downstream translation initiation candidates. First, we measured the occurrence of out-of-frame methionines downstream of SNVs, and found no enrichment for out-of-frame methionines downstream of nonsense variants relative to synonymous variants (**Figure S4B-D**). Second, we searched for in-frame methionines downstream of SNVs, but restricted to variants within the middle of the CDS, where downstream translation initiation would produce a severely truncated protein. We observed no statistically significant differences in occurrence of downstream methionines for synonymous versus nonsense variants (**Figure 4C**). We conclude that in-frame methionines are specifically enriched downstream of nonsense variants that lie near the 5’ ends of CDSs.

Finally, we tested whether nonsense variants that induced C-terminal truncation might be subject to relaxed selection. Approximately 10% of synonymous variants fell within the last 10% of their host CDSs, independent of allele frequency. In contrast, 13% of rare nonsense variants fell within this region, increasing to 21% for common nonsense variants (**Figure 4D**). Therefore, we conclude that the positional distribution of nonsense variants within CDSs is consistent with frequent induction of relatively small N- or C-terminal truncations of the encoded protein.

Ribosome profiling data similarly indicated that downstream translation initiation contributes to translation of mRNAs containing nonsense variants. We observed likely translation initiation downstream of nonsense variants at both canonical (ATG; *CCHCR1*) and non-canonical (CTG or GTG; *ABHD14B*) start codons (**Figure 4E-F, S4E-F**). Translation initiation at non-canonical start codons has been previously observed in mammalian genomes, although it is uncommon relative to initiation at canonical start codons (Ingolia et al. 2011). We confirmed that there are no regions in the reference human genome assembly with high homology to the exon containing the *ABHD14B* nonsense variant, and also observed 100% expression of the alternate allele in the 1/1 sample, indicating that genotyping errors are unlikely. Together, our data provides both sequence-based and direct evidence of translation initiation at canonical and non-canonical start codons downstream of common nonsense variants.

### Truncated proteins encoded by nonsense variant-containing mRNAs are produced in a heterologous system

Of the nonsense variants highlighted above (within *LGALS8*, *MOB3C*, *TMEM218*, *PVRIG*, *SLFN13*, *CCHCR1*, and *ABHD14B*), we could only confirm LGALS8 and MOB3C protein levels due to mass spectrometry’s incomplete coverage of the proteome. SILAC data was not available for proteins affected by the other highlighted nonsense variants. We therefore used a reporter system to experimentally confirm the conclusions of our genomic analyses, as well as test whether translation of nonsense variant-containing mRNAs could be recapitulated in a heterologous system (versus the endogenous context in which the genomic assays were performed). We focused on nonsense variants that we predicted to be subject to stop codon readthrough or induce N- or C-terminal protein truncation, as those involve incompletely understood biochemical mechanisms or the production of novel proteins not present in cells lacking the nonsense variant.

We designed a series of reporter constructs containing the reference (sense codon) or variant (nonsense codon) alleles of ten nonsense variants (**Table 1**). We chose to test variants within *ABHD14B*, *CCHCR1*, *PVRIG*, and *SLFN13* (**Figure 3–4**), as well as a blindly chosen set of nonsense variants that were present as 1/1 in the 1000 Genomes cohort but for which we did not have ribosome profiling data available for 1/1 samples. Each construct carried an N-terminal FLAG tag and a C-terminal HA tag (**Figure 5A**), permitting us to specifically identify the encoded protein and distinguish stop codon readthrough from N- or C-terminal truncation as follows. Upon translation, the reference allele is expected to produce a full-length protein product with both the N- and C-terminal tags. If the variant allele is subject to stop codon readthrough, then we expect to similarly observe a full-length product carrying both tags. In the case of N- or C-terminal truncation, however, only a partial protein product carrying a HA or FLAG tag, respectively, is expected (**Figure 5B**).

**Table 1.** Characteristics of nonsense variants selected for reporter assays. *q*, allele frequency of the alternate allele in the 1000 Genomes data. ASE, allele-specific expression based on RNA-seq. ASE was computed for each variant by restricting to samples that were heterozygous (0/1) for the variant and had at least ten reads covering the variant. No 0/1 samples had sufficient reads to compute ASE for the variant in *FCGR2A*. “Isoform-specific” was defined as in Figure 2A. “Relative pos. (%)” indicates the relative position of the codon containing the SNV within the host CDS. “Reference validated” indicates that the construct containing the reference allele produced a protein product of the expected length with both N-and C-terminal tags. RefSeq ID, ID of the RefSeq parent transcript for which each variant introduces an in-frame stop codon. Locus, genomic coordinates for the GRCh37/hg19 genome assembly. Alleles, nucleotides associated with the reference>alternate alleles.

|  | RefSeq ID | locus | alleles | *q*<br>(%) | ASE<br>(%) | isoform-specific | relative<br>pos. (%) | ref. val-<br>idated | origin of<br>protein |
| --- | --- | --- | --- | --- | --- | --- | --- | --- | --- |
| *ABHD14B* | NM_001146314 | 3:52,005,638 | G>A | 1.3 | 37 | yes | 7.6 | yes | no protein |
| *CCHCR1* | NM_001105564 | 6:31,124,849 | C>T | 47.0 | 43 | yes | 8.9 | yes | downstream<br>initiation |
| *CCHCR1* | NM_001105564 | 6:31,125,257 | C>A | 10.0 | 40 | yes | 4.6 | yes | downstream<br>initiation |
| *FCGR2A* | NM_001136219 | 1:161,476,204 | C>T | 9.6 | - | yes | 19.5 | no | - |
| *NDUFV3* | NM_021075 | 21:44,323,720 | C>T | 6.4 | 16 | yes | 42.0 | yes | C-terminal<br>truncation |
| *PVRIG* | NM_024070 | 7:99,817,585 | G>T | 24.0 | 35 | no | 5.2 | no | - |
| *SDR42E1* | NM_145168 | 16:82,033,810 | G>A | 13.0 | 50 | no | 7.4 | no | - |
| *SLFN13* | NM_144682 | 17:33,772,658 | G>T | 10.0 | 53 | yes | 1.6 | no | - |
| *TLR10* | NM_030956 | 4:38,776,104 | G>A | 1.4 | 45 | no | 45.4 | no | - |
| *TRIM38* | NM_006355 | 6:25,969,631 | C>T | 3.0 | 15 | no | 35.0 | yes | C-terminal<br>truncation |

**Figure 5.**
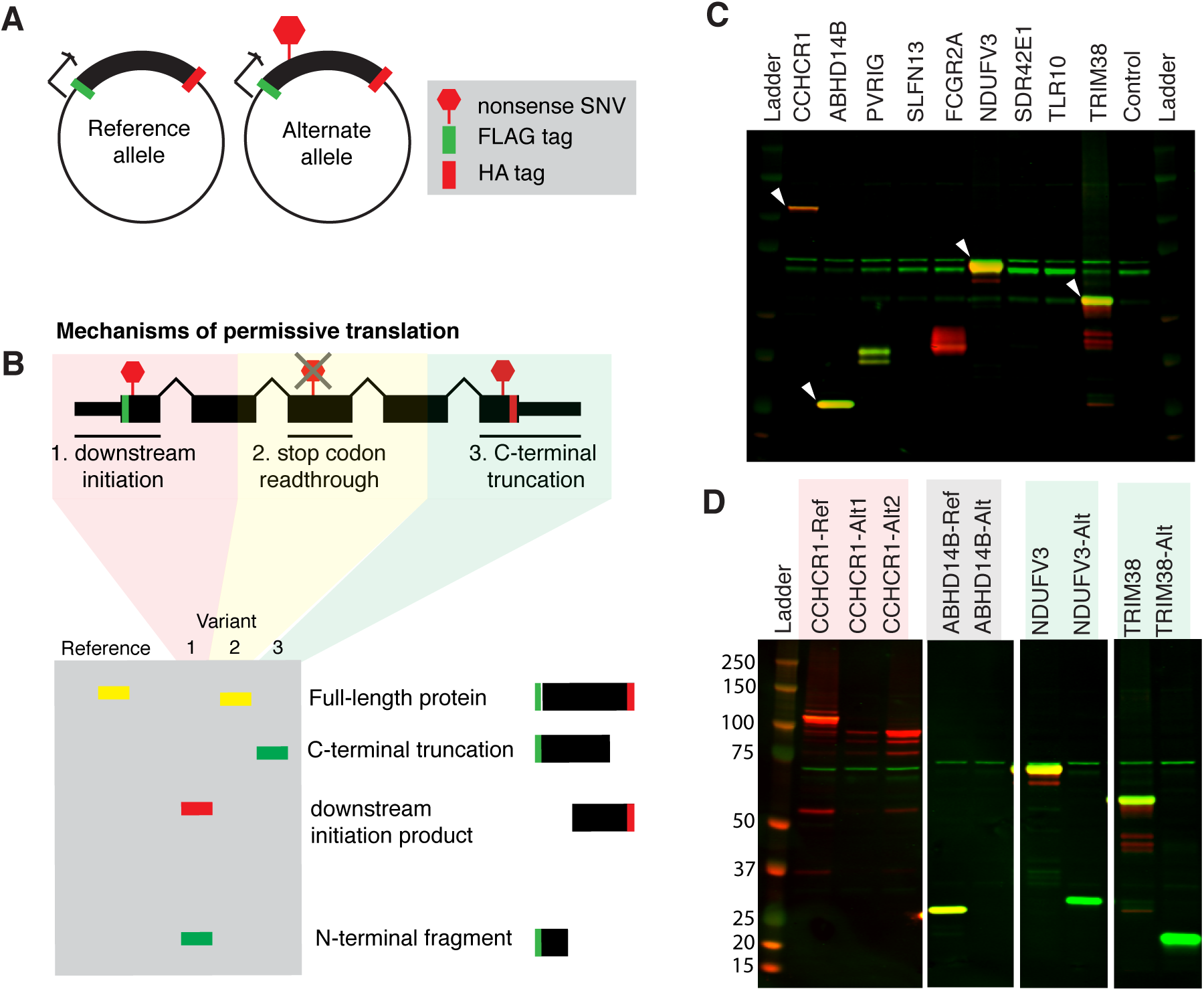
A reporter system recapitulates translation of N- and C-terminally truncated proteins from mRNAs containing nonsense variants. (**A**) Design of reporter constructs carrying the reference (sense) or alternate (nonsense) alleles of selected genes. (**B**) Cartoon depicting how permissive translation may enable productive translation of mRNAs containing nonsense variants, as well as the expected protein products from our reporter constructs for each mechanism of permissive translation. (**C**) Western blot of total protein from HEK293 cells transfected with constructs containing the reference alleles of all candidate genes. The FLAG and HA tags are illustrated in green and red. White arrows indicate constructs that produced the expected protein with both FLAG and HA tags. (**D**) As (C), but for constructs containing either the reference (Ref) or alternate (Alt) alleles of the genes marked with white arrows in (C). For CCHCR1, the “Alt1” and “Alt2” variants corresponds to SNVs at genomic positions 31,124,849 and 31,125,257 on chromosome 6. The FLAG and HA tags are illustrated in green and red.

We introduced each construct containing the reference or variant allele of each candidate gene individually into HEK293 cells via transient transfection and assayed their protein products 48 hours post-transfection by detecting the FLAG and HA tags. Constructs containing the reference alleles of *CCHCR1*, *ABHD14B*, *NDUFV3*, and *TRIM38* all produced a protein product of the expected size carrying both N-and C-terminal tags (**Figure 5C,S5A**). The other reference constructs either produced no protein or protein products that were not of the expected size, and so were excluded from further analysis. We additionally tested constructs for *PVRIG* and *SLFN13* that contained their endogenous 5’ and 3’ untranslated regions (UTRs), but found that those constructs also did not produce the expected protein products (data not shown). The absence of detectable protein from some constructs may be due a requirement for cell type-specific translational machinery that is absent from HEK293 cells, problems caused by our introduction of N- and/or C-terminal tags, or possibly incorrect coding sequence annotations. The encoded proteins might also be unstable in HEK293 cells.

Constructs containing either of the two nonsense variants of *CCHR1* produced an N-terminally truncated protein fragment indicative of initiation downstream of the premature termination codon (**Figure 5D,S5B**). This fragment was produced from the construct containing the reference allele as well, suggesting that *CCHCR1* is subject to alternative translation initiation even in the absence of a nonsense genetic variant. The construct containing the nonsense variant of *ABHD14B* did not make a detectable protein in HEK293 cells, despite our expectation given uninterrupted ribosome footprints throughout the open reading frame in LCLs, suggesting that an N-terminally truncated protein is likely translated but unstable. Constructs containing the *NDUFV3* and *TRIM38* nonsense variants produced C-terminally truncated protein fragments that were abundant despite their small size relative to the full-length proteins. These variants in *NDUFV3* and *TRIM38* were expressed in RNA-seq from 0/1 samples at allelic levels of 16% and 15% (**Table 1**), indicative of incomplete degradation by NMD that could potentially enable these short proteins to be similarly produced from their endogenous loci. In total, we detected stable expression of N- and C-terminally truncated protein products from constructs encoding four of five unambiguously tested nonsense variants.

### Stop codon readthrough rescues protein production from an endogenous gene containing a nonsense variant

As technical limitations of our heterologous reporter system prevented us from experimentally confirming that readthrough occurs for PVRIG and SLFN13 mRNAs, as predicted from ribosome profiling data, we turned to measurements of endogenous protein levels. We identified LCLs that were homozygous for either the reference (0/0) or nonsense (1/1) variants in *PVRIG* and *SLFN13* and obtained five such cell lines from the Coriell Biorepository (**Figure 6A**). We used an enrichment strategy to detect endogenous protein, wherein we immunoprecipitated using antibodies directed against PVRIG or SLFN13 and then probed the immunoprecipitate using the same antibodies (**Figure 6B**). We detected PVRIG protein at the expected molecular weight in LCLs that were homozygous for either the reference (0/0) or nonsense (1/1) variants, supporting our computational prediction that stop codon readthrough rescues protein production from the nonsense variant-containing allele of *PVRIG*. PVRIG protein was detectable in the LCLs after enrichment of the endogenous protein using two distinct antibodies, as well as in a positive control consisting of lysate from PVRIG-overexpressing cells, confirming the specificity of our assay (**Figure 6C-D**). SLFN13 was not detectable in any LCL, suggesting that either the two tested antibodies against SLFN13 are not effective probes for immunoblotting or that SLFN13 is an unstable protein. Together with data from our heterologous reporter system, these measurements of endogenous protein levels support our hypothesis that diverse mechanisms of permissive translation enable protein production from genes containing nonsense variants.

**Figure 6.**
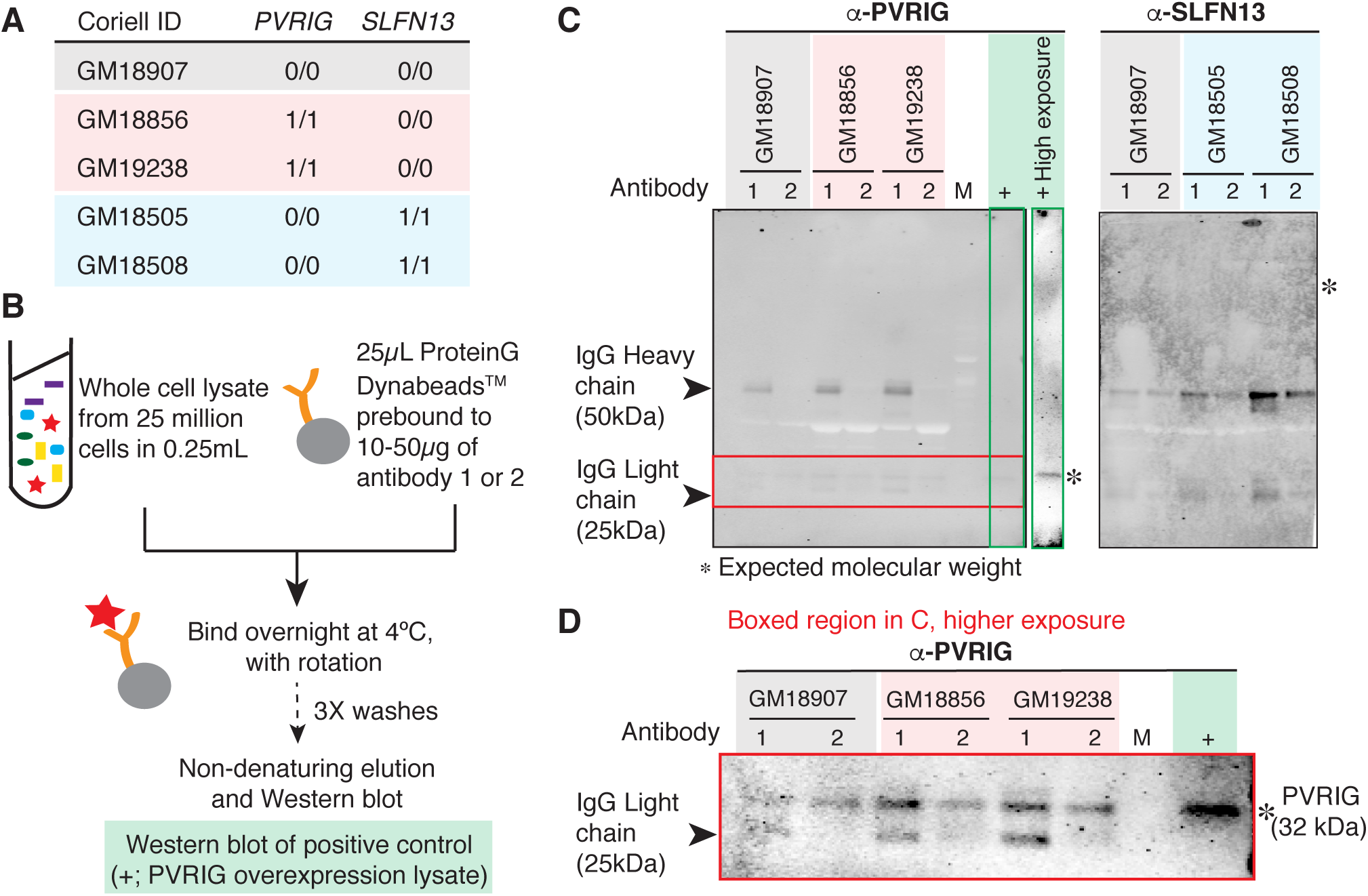
Stop codon readthrough enables protein production from a gene containing a nonsense variant. (**A**) Lymphoblastoid cell lines (LCLs) used in this study and their genotype for *PVRIG* and *SLFN13*. (**B**) Cartoon depicting the experimental setup for immunoprecipitation and subsequent Western blotting for PVRIG and SLFN13 to measure endogenous protein in the LCLs listed in (A). (**C**) Western blots for PVRIG and SLFN13 on immunoprecipitates that were enriched for the corresponding proteins from the illustrated LCLs. Two distinct antibodies were separately used for protein enrichment and the antibodies were used together for Western blot detection. *, expected molecular weight for PVRIG (detected by both antibodies) and SLFN13 (detected by neither antibody). M, marker lane containing the ladder. +, positive control for PVRIG (PVRIG overexpression lysate). + High exposure, lane for the positive control (green box) shown in a higher exposure to illustrate specificity of the assay. Note that even though non-denaturing elution limits the bulk release of the Protein G-bound antibodies that were used to enrich for the antigen, some antibody release is unavoidable and is visible as Heavy and Light chains, which migrate at 50kDa and 25kDa. Those bands do not interfere with the detection of PVRIG and SLFN13, which are expected to migrate at 32kDa and 102 kDa, respectively. (**D**) Higher exposure of the relevant portion (red box) of the PVRIG blot in (C). PVRIG was detected by both antibodies at the expected molecular weight in all cell lysates.

## DISCUSSION

Our data demonstrate that many common nonsense variants have only modest impacts upon the levels of total protein produced from their seemingly disabled parent genes. Translation of mRNAs transcribed from genes containing nonsense variants arises from diverse mechanisms during the gene expression process, including transcription start and stop site choice, alternative splicing, translation start site choice, downstream translation initiation, and stop codon readthrough. Given the severe fitness costs of many complete gene knockouts in murine models (Ayadi et al. 2012; de Angelis et al. 2015), our data therefore suggest that permissive RNA processing and translation in human cells facilitates the accumulation of otherwise deleterious genetic variation in the human population.

What fraction of nonsense variants disable gene function? We were able to directly answer this question for common variants by relying on individuals that are homozygous for the variants of interest. However, a direct answer is inaccessible for rare variants, which almost never occur in a homozygous state. (The low coverage of ribosome profiling and SILAC data prevents reliable analysis of allele-specific translation or protein levels in heterozygotes.) We therefore used sequence analyses to estimate that 43-49% of rare and low-frequency variants identified by the 1000 Genomes Project are isoform-specific or may induce N- or C-terminal truncation (**Table 2**). These predictions do not take into account the incompleteness of current isoform annotations, the possibility of initiation at non-canonical start codons, or the possibility of stop codon readthrough, and so may be underestimates. We therefore speculate that permissive RNA processing frequently rescues protein production from genes containing rare nonsense variants, although at lower rates than their common counterparts.

**Table 2.** Percentages of nonsense variants that are predicted to potentially not abrogate protein production. Isoform-specific variants are defined as in Figure 2A. Candidates for downstream initiation and C-terminal truncation are defined as in Figure 4. The last row (total) specifies the percentage of variants that are isoform-specific, are candidates for downstream initiation, or cause C-terminal truncation. As described in the Discussion, these numbers are likely underestimates, and do not include the possibility of stop codon readthrough.

|  | rare | low | common |
| --- | --- | --- | --- |
| isoform-specific | 30.2% | 36.9% | 40.2% |
| downstream initiation | 5.5% | 5.7% | 9.8% |
| C-terminal truncation | 13.4% | 18.4% | 21.4% |
| total | 42.8% | 48.6% | 52.7% |

It is important to note that maintenance of protein production may not be equivalent to rescue of gene function in many cases. Different isoforms of a gene may encode proteins with distinct or even opposing functions, so that an isoform-specific nonsense variant may have unpredictable consequences. Even modest N- or C-terminal truncation could alter or abolish protein function by removing peptide signals that direct subcellular localization, prenylation, or other activities or modifications, or induce unexpected gain of function by placing a normally internal peptide signal at the N or C terminus. For example, a recent study identified a novel C-terminal truncating variant in *CD164* that segregated with nonsyndromic hearing impairment in an affected family. Only the last six amino acids of CD164 were removed, but without the encoded canonical sorting motif, the truncated protein was trapped on the cell membrane (Nyegaard et al. 2015).

The most unexpected finding of our study was the prominent role played by mRNA translational plasticity in enabling protein production from genes containing non-pathogenic nonsense variants. The regulation of mRNA translation initiation and termination is incompletely understood, yet relevant for interpreting genetic data. Our data indicate that downstream translation initiation is associated with relaxed purifying selection and gives rise to N-terminally truncated proteins for some nonsense variants. Our data also show that stop codon readthrough enables translation of at least one mRNA containing a common nonsense variant. Stop codon readthrough is thought to be relatively rare in mammals, with only a few known examples (Schueren et al. 2014; Stiebler et al. 2014; Loughran et al. 2014; Eswarappa et al. 2014). Nonetheless, efficient readthrough of an otherwise fatal nonsense variant has been observed clinically (Pacho et al. 2011), highlighting the importance of translational plasticity when interpreting genetic data.

Together, our data demonstrate that permissive RNA processing has the potential to convert loss-of-function variants from genetic nulls into hypomorphic, silent, or even neomorphic alleles. Our results may therefore help to explain recent observations that some apparently healthy individuals carry homozygous putative loss-of-function variants within genes implicated in Mendelian disease (Kaiser et al. 2015). An improved mechanistic understanding of RNA processing and translation is essential for ongoing attempts to predict the pathogenicity of genetic variation (Kircher et al. 2014; MacArthur et al. 2014).

## METHODS

### Genotypes

Genotypes from the phase 3 variant set called by the 1000 Genomes Project (1000 Genomes Project Consortium 2012) were downloaded from here: ftp://ftp-trace.ncbi.nih.gov/1000genomes/ftp/release/20130502 We restricted our analysis of ribosome profiling data from Battle et al (Battle et al. 2015) to the 62 samples with genotypes called in this variant set, and used the 1000 Genomes genotyping for all analyses. For the RNA-seq data from Lappalainen et al (Lappalainen et al. 2013), we used the genotyping information from the original study, available here: http://www.ebi.ac.uk/arrayexpress/files/E-GEUV-1/genotypes/ and restricted to the 421 samples that were genotyped by sequencing. For global SNV analyses based on the 1000 Genomes variant set, we restricted to SNVs for which the alternate allele was present in a subsampled fraction of the 1000 Genomes data, wherein we restricted each of the 26 populations to no more than 90 samples to help to even the genotyping coverage among the available populations. For all datasets, we restricted genotyping calls to SNVs.

### SNV functional annotation

We used SnpEff (Cingolani et al. 2012) to identify variants that affected coding DNA sequences and annotate their functional effects. We ran snpEff v4_1c with the options ‘-v -lof -no-downstream -no-intergenic -no-intron -no-upstream -no-utr --formatEff --download hg19’, and then restricted to SNVs with predicted effects of ‘synonymous_variant’ or ‘stop_gained’ for at least one RefSeq transcript. For all transcript-level analyses, such as computing the amino acid affected by a SNV, we relied upon the AnnotationHub and BSgenome packages within Bioconductor (Huber et al. 2015).

### Genome annotations

A merged genome annotation was created by combining the UCSC knownGene annotations (Meyer et al. 2013), Ensembl 71 (Flicek et al. 2013), and isoform annotations from MISO v2.0 (Katz et al. 2010) as previously described (Dvinge et al. 2014). A file of all possible splice junctions, consisting of all splice junctions present in the merged genome annotation as well as all possible combinations of 5’ and 3’ splice sites within each parent gene, was created for subsequent read mapping.

### RNA-seq and ribosome profiling read mapping

FASTQ files were downloaded for RNA-seq data from LCLs from EBI’s ArrayExpress under accession number E-GEUV-1 (Lappalainen et al. 2013), for ribosome profiling data from LCLs from the NCBI Gene Expression Omnibus under accession number GSE61742 (Battle et al. 2015), and for the Illumina Body Map 2.0 dataset from ArrayExpress under accession number E-MTAB-513. RNA-seq and ribosome profiling reads were mapped to the UCSC hg19 (NCBI GRCh37) genome assembly as previously described (Dvinge et al. 2014). (Because our analyses consisted of comparisons between the relative expression of the reference versus alternate alleles, which are unaffected by the specific genome assembly, using the NCBI GRCh38 genome assembly instead of GRCh37 would not significantly affect our conclusions.) Briefly, RSEM (Li and Dewey 2011) was used to map reads to annotated transcripts, and the resulting unaligned reads were then aligned to the genome and the set of all possible splice junctions using TopHat v2.0.8b (Trapnell et al. 2009). We required that reads map with a maximum of three mismatches for the RNA-seq data, which consisted of 2x75 bp reads, and one mismatch for the ribosome profiling data, which consisted of shorter mixed-lengths reads. Reads mapped by RSEM and TopHat were then merged to create final BAM files of aligned reads.

### SILAC mass spectrometry data

Gene-level quantification of relative protein abundance based on SILAC was taken from Supplementary Data Table 4 of Battle et al (Battle et al. 2015).

### Transcript expression, allele-specific expression, and isoform ratio measurements

Transcript expression (e.g., as illustrated in **Figure 1C-D**) was quantified for RNA-seq or ribosome profiling data in units of fragments per kilobase per million mapped reads as follows. For each coding gene annotated in Ensembl 71, we counted the numbers of reads (or fragments, in the case of paired-end reads) that aligned to the longest CDS of that gene based on the RefSeq transcript set and then divided by the CDS length in kilobases. We then normalized these counts by dividing by a sample-specific normalization factor, defined as (10^6^ × total number of reads mapping to any Ensembl coding gene as specified above). This computation relied upon the GenomicAlignments, GenomicFeatures, and GenomicRanges packages within Bioconductor (Lawrence et al. 2013; Huber et al. 2015), and resulted in expression measurements in units of fragments per kilobase per million mapped reads (FPKM).

Allele-specific expression was measured by counting the numbers of RNA-seq reads containing the reference or alternate alleles. Isoform ratios were quantified genome-wide using MISO (Katz et al. 2010) as previously described (Dvinge et al. 2014).

### Read coverage plots

Ribosome profiling read coverage plots were generated from BAM files of mapped reads as follows. For each sample, all reads aligned to the genomic locus of interest were extracted and trimmed to 25 bp by removing equal lengths of sequence as necessary from the 5’ and 3’ ends to control for the differing footprint read lengths. At each genomic position, the normalized coverage was computed as the total number of reads aligned to that position divided by a sample-specific normalization factor, defined as (10^6^ x total number of reads mapping to any Ensembl coding gene). Samples were then classified according to their genotype and the average coverage was computed for each genotype by averaging over the normalized coverage values for each sample. RNA-seq read coverage plots were generated via an identical procedure, but the reads were not trimmed (as all reads were of equal length). These plots relied upon the AnnotationHub and BSgenome packages within Bioconductor (Huber et al. 2015).

### Plotting and graphics

All plots and figures were generated with the dplyr (Wickham and Francois) and ggplot2 packages (Wickham 2009).

### Reporter constructs

The reporter constructs for reference alleles were obtained as full-length cDNAs in the pcDNA3.1 backbone carrying FLAG and HA tags at the N and C termini, respectively (GenScript USA Inc.). Point mutations were introduced into constructs containing the reference alleles to generate the variants (GenScript USA Inc.).

### Immunoprecipitation of PVRIG and SLFN13 from lymphoblastoid cell lysates

Lymphoblastoid cell lines were obtained from the Coriell Biorepository and cultured under recommended conditions in RPMI with 15% FBS. 25 million cells were pelleted and lysed in NP-40 lysis buffer (50mM Tris, pH 8.0, 150mM NaCl, 1% NP-40) containing EDTA-free protease inhibitor mini tablets (Pierce; Cat. # 88666). The lysate was incubated overnight at 4°C with 25µL of protein G Dynabeads (Invitrogen) prebound to 10-50µg of antibody (PVRIG: ab168153 (Abcam Inc.), NBP2-13832 (Novus Biologicals); SLFN13: ab173951 (Abcam Inc.), NBP1-93879 (Novus Biologicals). The beads were washed thrice with the Dynabead wash buffer and the bound proteins eluted in the non-denaturing elution buffer. The eluate was then subjected to Western blotting as described below using the same antibodies used for the immunoprecipitation. A transient overexpression lysate of PVRIG (LY411386; Origene Technologies, Inc.) served as a positive control to determine the specificity of the antibodies used for Western blotting.

### Western blotting

HEK293 cells were grown in DMEM containing 10% FBS and transfected with the reporter constructs using Lipofectamine^®^ 3000 transfection reagent (Invitrogen). After 48 hours, the cells were lysed in TRIzol^®^ reagent (Invitrogen). Total protein was extracted following the manufacturer’s instructions the protein pellet resuspended in sample buffer containing 5%SDS and 0.5M unbuffered Tris base to ensure efficient solubilization, aided by gentle sonication in an ultrasonic water bath. Protein concentrations were determined using the BCA protein assay (ThermoScientific) and 5 µg of total protein was resolved on 4-12% NuPAGE Bis-Tris acrylamide gels (Novex) and western blotting was performed using the LICOR system. The primary antibodies used were mouse monoclonal anti-FLAG M2 antibody (Sigma-Aldrich; Cat # F1804) and rabbit monoclonal anti-HA antibody (EMD Millipore; Cat # 05-902R). IRDye-conjugated secondary antibodies (LICOR) were used for quantitative detection of the tags. Rough estimates of the protein molecular weights were made by comparison with the Precision Plus Protein Kaleidoscope standards (BioRad).

## ACKNOWLEDGEMENTS

This research was supported by the Ellison Medical Foundation AG-NS-1030-13 (RKB) and the FSH Society FSHS-22014-01 (SJ). We thank Jesse Bloom and Guo-Liang Chew for comments on the manuscript.

## AUTHOR CONTRIBUTIONS

SJ and RKB performed the experiments, analyzed the data, and wrote the paper.

## DISCLOSURE DECLARATION

The authors declare that no competing interests exist.

